# Progeny differentiation in faba bean using hyperspectral images and machine learning

**DOI:** 10.64898/2026.05.19.725957

**Authors:** Rica-Hanna Schlichtermann, Sebastian Warnemünde, Hanna Tietgen, Gregor Welna, Andreas Stahl, Benjamin Wittkop, Rod J. Snowdon

## Abstract

Though currently a minor crop, faba bean is a promising source of plant-based protein as global diets shift towards more plant-based nutrition. To realise this potential, advances in breeding and cultivation are crucial. To exploit heterosis, faba bean breeding frequently utilises synthetic cultivars, which involves open pollination of inbred lines to produce a mixture of F_1_ hybrid seeds and self-pollinated offspring. Pure F_1_ hybrid cultivars are currently unavailable due to unstable cytoplasmic male sterility (CMS) systems. An ability to distinguish F_1_ seeds from their parental inbreds via characteristics associated with xenia effects could change this. The xenia effect refers to the influence of paternal pollen on seed traits, for example seed weight and cotyledon cells in faba bean. In this study, we exploited the xenia effect captured in hyperspectral imaging data to develop machine learning scenarios for discriminating between parental and F_1_ seeds of open pollinated synthetic combinations (Syn-1). The hyperspectral data were pre-processed using Savitzky–Golay filtering to reduce noise and smooth the spectra. Various machine learning algorithms were applied, incorporating Bayesian hyperparameter optimisation. The scenarios achieved up to 98.9 % accuracy in separating parental components of Syn-1. When including all seeds, the model achieved 40.7 %, indicating moderate detection and classification performance. As the harmonic mean of precision and recall, the F1 score accounts for both the correctness of F_1_ seed detections and the completeness with which F_1_ seeds were detected. While this approach does not yet enable the development of full hybrid cultivars, it paves the way for hybrid-enriched cultivars. These could help to streamline breeding for synthetic cultivars and potentially increase yields, for example by increasing the proportion of F_1_ hybrid seeds in synthetic cultivars. This study extends knowledge of the xenia effect in faba bean and provides a basis for further research aimed at enhancing breeding methods and productivity.

## 1 Introduction

With an increasing global population and a rising demand for plant-based protein, production of faba bean (*Vicia faba* L.) has expanded, underscoring its growing importance (Food and Agriculture Organization 2022, Sim et al. 2021). Faba bean offers significant benefits, such as a relatively strong ability to fix atmospheric nitrogen in symbiosis with *Rhizobium* bacteria, thereby providing residual nitrogen for subsequent crops (Aschi et al. 2017, Liu et al. 2019). This advantage is evident in improved yields of up to 20 % in crop rotation systems that incorporate legumes compared to non-legume systems (Zhao et al. 2022a). Additionally, faba bean seeds are characterized by a high protein and mineral content, making them suitable for use in alternative protein products, such as meat substitutes (Augustin and Cole 2022). Despite these benefits, faba bean suffers from yield instability, primarily due to drought and heat stress, necessitating further research, particularly in the breeding process, to fully harness its potential (Food and Agriculture Organization 2022, Bishop et al. 2016, Balko et al. 2023, Lavania et al. 2015).

Faba bean shows significant levels of heterosis (Stelling 1997, Link et al. 1996), despite cytoplasmic male sterility systems (CMS) in faba bean being found (Bond et al. 1964, Palmer et al. 2011), these are note reliable so that breeding of pure F_1_ hybrids is presently not possible. Instead, to at least partially exploit heterosis, faba bean is frequently bred as synthetic cultivars (Link 1990). This breeding process involves combinations of at least two selected inbred lines, which are propagated together in the field as a Syn-0 combination. Open pollination and the allogamous nature of faba bean then facilitates outcrossing, resulting in a mixture of harvested seeds from F_1_ seed combinations between the inbred lines along with self-pollinated inbred seeds. Subsequent field propagation again under open pollination of this first synthetic generation (Syn-1) results in a Syn-2 generation comprising a population of seeds from self-pollinated parental genotypes, F_2_ progenies, parent x F_1_ crosses and (in case more than two founder parents were used) multi-cross hybrids (F_1_ x F_1_). Due to the heterozygous state, these synthetic generations tend to exhibit increased and more stable yields, showcasing the heterosis effect. Highest yields are often achieved in Syn-2 and Syn-3 generations, depending on the number of parental components and their degree of self- and cross-pollination, hence up to two generations of propagation under open pollination may be performed (Stelling et al. 1994). This adds considerable time and expense to breeding operations. A considerably more efficient scenario to fully harness heterosis in faba bean, which can range from 33 % to 51 % for mid-parent yield heterosis (Link et al. 1994b), would be to breed F_1_ hybrids (Melchinger and Gumber 1998). However, in the absence of an effective CMS system, alternative strategies for exploitation of heterosis in the F_1_ generation might be advantageous to overcome the cost and effort associated with breeding of advanced-generation synthetic cultivars.

In 1881, Focke described how fruits from crosses between *Raphanus sativus* and *Raphanus raphanistrum* could be distinguished from the two parents based on their colour. He described the possible influence of different factors carried by the pollen, indicating that hybridity could affect fruit characteristics. Focke (1881) termed this phenomenon “xenien”. Today, the term “xenien” or “xenia” is used to describe the general influence of the pollen on fruit development, including colour, shape, size, and chemical components (Denney 1992). Well-known examples of xenia effects include increased seed weight in maize, and effects on colour and shape in offspring of inter- and intraspecific crosses in peas (Bulant and Gallais 1998, Sari et al. 2023). In faba bean, an increased number of cotyledon cells and seed weight have been observed (Duc et al. 2001, Dieckmann and Link 2010). These effects, reflecting the influence of paternal pollen on the seed, can be utilized, as proposed by Sari et al. (2023), to expedite crossing processes and reduce costs through reduced genotyping.

Spectral data has been used for quality assessment in crop production and breeding since the 1960s, when near-infrared reflectance (NIR) patterns were first shown to be associated with seed moisture content (Hart et al. 1962). More recently, it was demonstrated that hyperspectral images and machine learning could be used for seed variety identification in wheat, okra and loofah seeds, with accuracy ranging between 95 to 98 % (Zhao et al. 2022b, Nie et al. 2019). Extending the use of variation in seed NIR spectral patterns, Rincent et al. (2018) introduced the concept of phenomic selection. Roscher-Ehrig et al. (2024) suggested that, especially in the start of a breeding programme, phenomic selection on the basis of NIR data could help to select superior genotypes, leading to better prediction accuracies with lower costs. New hyperspectral imaging technologies extend the wavelength range of reflectance measurements beyond the infra-red and visual spectra. Here, we propose that linking of hyperspectral imaging to the xenia effect in faba bean and other crops could enable a high-throughput phenotyping approach that might facilitate the creation of F_1_ hybrid cultivars, or at least hybrid-enriched synthetic cultivars. This can potentially help to implement the benefits of hybrid breeding, including improved yield and yield stability (Lee and Tollenaar 2007, Huang et al. 2017, Hackauf et al. 2022), into crops such as legumes for which establishment of traditional hybrid breeding systems based on biotechnological fertility control mechanisms has to date been unsuccessful.

This study investigates the use of hyperspectral imaging in combination with machine learning to differentiate the parental components of biparental Syn-1 seed combinations. The approach is further extended to distinguish F_1_ hybrid seeds from homozygous selfed seeds in Syn-1 combinations, in order to analyse whether an observed xenia effect can be detected in this context. This could lead to potential applications in plant breeding to develop Syn-1 cultivars enriched for a higher proportion of F_1_ hybrid seeds, either through the accurate identification of F_1_ seeds or through classification and use of parental components that exhibit a higher outcrossing rate. In a more general context, it could also help breeders reduce costs for genotyping to confirm hybridity of offspring from controlled crosses.

## 2 Material and Methods

### 2.1 Plant material

A total of 18 biparental, synthetic (Syn-1), spring-type faba bean mixtures, along with pure-breeding seeds from their respective parental lines, were obtained from the NPZ Innovation GmbH (NPZi) (Holtsee, Germany) from the BreedPath project. In this study, each Syn-1 combination comprised seeds harvested from two parental components after open pollination of two more or less homozygous parental lines under isolation tents with bumblebees to guarantee outcrossing, meaning that the Syn-1 comprises a mixture of F_1_ seeds from cross-pollination and inbred seeds from self-pollinations of the respective parental lines. Initially, 96 seeds from each Syn-1 were analysed along with 12 seeds from each parental line, and phenotypic data were collected for seed weight. Seeds were transferred to 24-well tissue culture trays for hyperspectral imaging, for each individual tray, as described below in detail. Individual seed images were extracted from the tray images using segmentation algorithms. Subsequently, the same procedure was performed using 300 additional seeds from two selected Syn-1 combinations (NPZ-BP3Y2023-007 & NPZ-BP3Y2023-060, respectively). All spectral-imaged seeds were grown for DNA extraction and marker analysis to determine whether their genotype corresponds to the F_1_ hybrid or one of the respective homozygous parents.

### 2.2 Overview of performed analysis

In this study the experiment was divided into two stages. Stage 1 focuses on a global scenario including all 18 Syn-1 combinations, with 96 seeds sampled from each Syn-1 combination (scenario 1). First utilizing this global scenario the parental components, the self-pollinated plants, were classified excluding the F_1_ seeds from the prediction. Furthermore, the classification capacity of the scenarios was assessed by investigating whether the F_1_ seeds could also be discriminated. To improve the prediction of F_1_ seeds, the F1 score and accuracy, different strategies were further used to investigate the full potential of this scenario. Previous studies showed that the xenia effect is more pronounced in more divergent genotypes (Duc et al. 2001). Hence, the Syn-1 combinations were ranked according to the degree of spectral divergence between their respective parental components, in order to identify genotype-specific differences. In Stage 2, Syn-1 combinations with a high outcrossing rate and whose parental lines were categorised as belonging to more “distant” spectral groups were chosen to be sampled again with a higher number of seeds (300 more seeds per Syn-1 combination, scenarios 2 and 3). Here, besides the parental differentiation and simple F_1_ prediction, a wavelength-specific approach was also applied to only utilize spectral patterns where the F_1_ differentiate from lying between the parental seeds. For scenario 3, the ability to distinguish F_1_ seeds using the spectral data was further investigated, separating the parental components of the Syn-1 based on a principal component analysis (PCA) (Table 1).

**Table 1:** Overview of used scenarios with target prediction, used data and applied strategies.

| Stage | Scenario | Prediction | Data structure | Applied strategies |
| --- | --- | --- | --- | --- |
| 1 | Scenario 1:<br>18 synthetic Syn-1 combinations with 96 seeds each | Parental | Balanced | Simple |
| | | $F_1$ | resampling of crosses | Simple, Distance groups |
| 2 | Scenario 2<br>NPZ-BP3Y2023-060 with 300 additional seeds | Parental | Balanced | Simple |
| | | $F_1$ | Balanced | Simple, specific wavelengths |
|  | Scenario 3<br>NPZ-BP3Y2023-007 with 300 additional seeds | Parental | Balanced | Simple |
| | | $F_1$ | Balanced | Simple, specific wavelengths, separated parental components |

### 2.3 Hyperspectral Imaging

For hyperspectral imaging, seeds were placed in 24-well tissue culture plates before sowing, with each seed being assigned a unique identifier. Hyperspectral imaging was conducted over a spectral range of 400 to 2500 nm. For this purpose, two hyperspectral line-scanning sensors (Norsk Elektro Optikk A/S, Skedsmokorset, Norway) were used: (i) the HySpex VNIR-1800, covering the visible and near-infrared region (VNIR; 400–1000 nm), and (ii) the HySpex SWIR-384, covering the short-wave infrared region (SWIR; 1000–2500 nm). Image acquisition and radiometric calibration were performed using the camera vendor’s software, HySpex Ground and HySpex Rad. In every image, a Zenith Polymer® diffuser (Sphere Optics GmbH, Herrsching, Germany) was included for calibration. The camera frame period was adjusted by the image acquisition software to match the speed of the linear stage, thereby ensuring geometrically correct image acquisition. For illumination a 1000 W short-wave halogen spotlight (Hedler C12, Hedler Systemlicht, Runkel/Lahn, Germany) was installed. The set up is shown in Figure 1.

**Figure 1:**
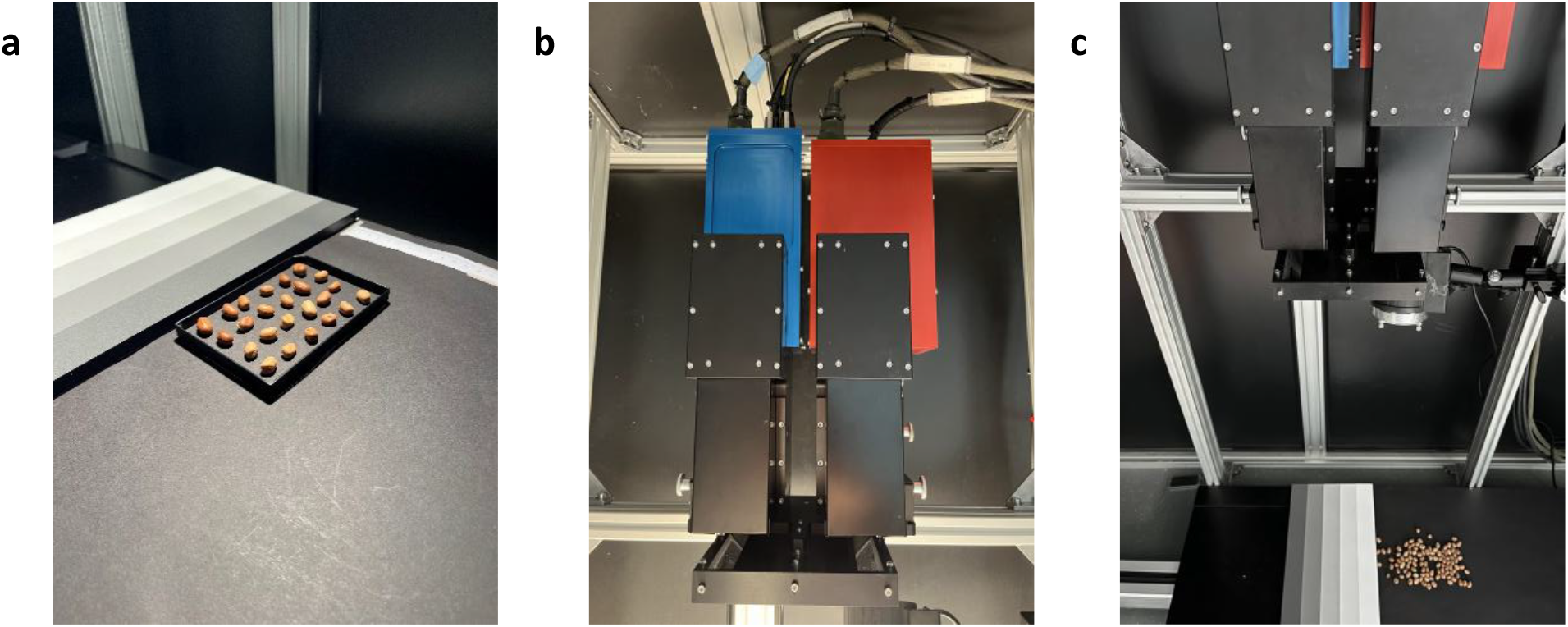
Set up for hyperspectral imaging. (a) Positioning of seeds (b) hyperspectral cameras (c) overview set up

Each measurement was calibrated using the Zenith Polymer® diffuser (white target) with known reflectance and the reflectance of the target was measured by the following equation (1)

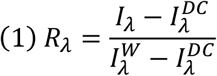

Where *I* _λ_ is the image pixel intensity at wavelength λ, 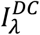 is the so called “dark current”, which is the intensity when measured with closed shutter and 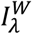, the intensity recording the white target. To remove the background, a segmentation scenario for the seed material was developed using the software perClass Mira (perClass BV, Delft, The Netherlands). For this purpose, a subset of images was manually annotated, and a reflectance-based scenario was trained to classify each spectral pixel as either seed or background. For each seed the mean reflectance, the reflectance standard deviation and the seed size were extracted from the images using perClass Mira object feature selection.

### 2.4 Leaf samples and DNA extraction

Leaf samples from plants grown under field or greenhouse conditions were collected in 1.1 mL deep well tubes on dry ice with 10 leaf discs of a diameter of 5 mm. Samples were freeze-dried for 2 days and extracted with the BioSprint 96 DNA Plant Kit from Qiagen (Düsseldorf, Germany) according to the manufacturer’s instructions. The DNA concentration of 5 samples was measured utilizing Qubit dsDNA BR Assay Kit from Invitrogen (Thermo Fisher Scientific, Waltham, MA, USA) as a reference and a standard dilution to 40 ng/µL was performed.

### 2.5 Syn-1 classification

Based on SNP chip analysis of the parental lines conducted by SGS Fresenius Institute GmbH TraitGenetics Section (Gatersleben, Germany), Kompetitive Allele Specific PCR (KASP) markers were designed to distinguish between parental components of Syn-1 combinations. For each Syn-1 combination, one marker was developed that could classify its offspring as homozygous (corresponding to one of its parental components) or heterozygous (F_1_-hybrid of the respective parents). KASP marker design was performed according to Makhoul and Obermeier (2022) and analysis followed the PCR programme in Supplementary Table S1. To confirm the functionality of the KASP markers, DNA samples from the respective homozygous parents were added to each KASP analysis as a control. Data was analysed using the QuantStudio™ Real-Time PCR software from applied biosystems® by Thermo Fisher Scientific (Waltham, MA, USA).

### 2.6 Machine learning

The following algorithm and R packages were utilized to distinguish the homozygous parental components from the heterozygous F_1_ seeds from the Syn-1 combinations: k-nearest Neighbour (KNN) (r-package “gmodels” (Warnes et al. 2024)), naive bayes (NB) (r package “e1071” (Meyer et al. 2024)), decision tree with the algorithm C5.0 (DT) (r-package “C50” (Kuhn and Quinlan 2025)), random forest (RF) (r-package “randomForest” (Liaw and Wiener 2002)) and support vector machine (SVM) (r-package “kernlab” (Karatzoglou et al. 2004)). We performed a 5-fold cross validation repeated 10 times with different set folds.

Evaluation of machine learning algorithms was based on the evaluation metrics precision (2), recall (3), F1 score (4), and accuracy (5), which were constructed based on the confusion matrix. The metrics were calculated using the following formulas:

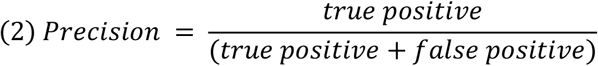

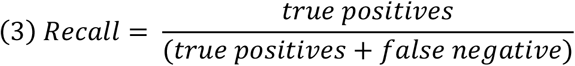

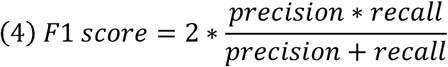

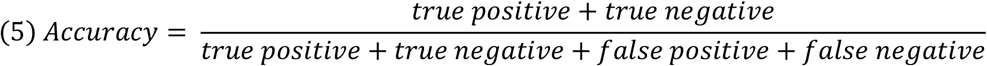

For differentiation of parental components based on spectral patterns, scenario performance during training was optimized using accuracy as the target metric. In contrast, for differentiation of the whole Syn-1, with a focus on F_1_ seeds, the F1 score was used as the optimization metric during training, as it provides a balanced measure of F_1_ hybrid seed identification and correct classification.

During preprocessing, the spectral data was filtered to reduce noise with Savitzky-Golay filtering, utilizing the R package “prospectr” (Stevens and Ramirez-Lopez 2025). The hyperparameters for the Savitzky-Golay filtering (windowsize, polynom and derivate) of each algorithm were tuned individually utilizing a grid search. The according hyperparameters are described in Table S2. Hyperparameters of the algorithm were tuned individually utilizing bayesian hyperparameter optimization (r-package “ParBayesianOptimization” (Wilson 2022)).

In the two stages, different approaches were taken to manage unbalanced numbers of samples in each class. In scenario 1, for the differentiation of parental components based on the spectral patterns, the number of samples were balanced through reducing the number of samples to the count of a single parent. For differentiation between all components of Syn-1 combinations, with the focus on identifying F_1_ seeds, adjustment of sample numbers by oversampling was necessary as the algorithm tended to overlook F_1_ seeds represented by only a small number of individuals. In scenario 1, the samples for each cross were duplicated in the training set to increase their representation. Due to the higher number of individuals in scenarios 2 and 3, a balanced dataset was used based on the number of parents in the differentiation of parental components of Syn-1 and the number of F_1_ seeds in the differentiation in the Syn-1 combinations. Syn-1 combination NPZ-BP3Y2023-060 consists of 82 F_1_ seeds, whereas Syn-1 combination NPZ-BP3Y2023-007 consists of 70 F_1_ seeds.

In scenario 1, the differentiation between the different Syn-1 combinations was coded as dummy variables, constructed with the R package psych (Revelle 2025). This is described as parental information (P). In all scenarios we tested the influence of the seed weight and seed size data, analysed and labelled together as (W). An approximation of the seed size was estimated by multiplying the seed length by the seed width. In combination with the spectral data, we tested scenario 1 with 4 datasets consisting of (1) spectral, weight, size and parental information (SWP), (2) spectral, seed weight and seed size (SW), (3) spectral and parental information (SP), and (4) only spectral data (S). In the scenarios 2 and 3, in which only one Syn-1 combination per scenario was used, only two datasets were necessary, consisting of (2) spectral, weight and size data (SW) and (4) only spectral data (S), as the parental information would have been the same for each entry.

In stage 1, the 18 Syn-1 combinations were divided into two groups based on the differences between the spectral data of their respective parental components, in order to investigate the influence of spectral differences between seeds of the respective parental components on the performance of the algorithms. The mean euclidian distance of the spectral data was calculated between parental component 1 and parental component 2 of each respective Syn-1 combination. The Syn-1 combinations were ranked from most distant to least distant and split into two equal-sized groups representing “distant” and “near” Syn-1 combinations (Table 2). The algorithms were run for each group separately and tuned individually.

**Table 2:** Assignment of Syn-1 combinations into “distant” and “near” Syn-1 combinations based on the mean euclidian distance of the spectra of their respective parental components.

| Distant | Near |
| --- | --- |
| NPZ-BP3Y2023-056 | NPZ-BP3Y2023-107 |
| NPZ-BP3Y2023-090 | NPZ-BP3Y2023-113 |
| NPZ-BP3Y2023-093 | NPZ-BP3Y2023-054 |
| NPZ-BP3Y2023-007 | NPZ-BP3Y2023-001 |
| NPZ-BP3Y2023-053 | NPZ-BP3Y2023-023 |
| NPZ-BP3Y2023-006 | NPZ-BP3Y2023-020 |
| NPZ-BP3Y2023-065 | NPZ-BP3Y2023-021 |
| NPZ-BP3Y2023-047 | NPZ-BP3Y2023-116 |
| NPZ-BP3Y2023-060 | NPZ-BP3Y2023-036 |

### 2.7 Principle Component Analysis

To visualize the spectral data, PCA was performed based on the data from scenario 1 for all Syn-1 combinations, based on modified Rogers’ genetic distance (Goodman and Stuber 1983, Wright 1978). For scenario 2 and 3, additional PCAs were performed including all samples of the respective Syn-1 combinations.

## 3 Results

### 3.1 Visualisation of spectral data

Visualization of Syn-1 combinations by PCA revealed distinct Syn-1 combination-specific differences. When the spectra from parental lines formed two separate PCA clusters, spectra from seeds of their respective F_1_ seeds were generally positioned within these clusters, indicating a strong maternal effect on the spectral signature of the seeds (which also encompasses the seed colour). Some Syn-1 combinations, however, did not exhibit clear cluster separation or display overlapping clusters (Supplementary Figure S1). Maternal effects were further evident in line graphs showing the mean spectra of the parental components and their F_1_ seeds for each Syn-1 combination. The parental lines form two distinct spectral profiles, with the F_1_ seeds spectra typically lying between them, building the mean spectra of the two parental components of each Syn-1 combination. Occasionally, F_1_ seeds exhibit higher or lower excitation at specific wavelengths, suggesting the presence of F_1_ seed-specific spectral characteristics (Supplementary Figure S2).

### 3.2 Stage 1

#### 3.2.1 Data structure

In our dataset, the percentage of F_1_ seeds among the Syn-1 combinations ranged from 13.5 % to 26.9 %, with a mean of 17.9 % among the F_1_ seeds analysed. The proportion of homozygous (self-pollinated) seeds, the parental components ranged from 11.8 % to 66.7 %, with a mean of 41 % per parental component per Syn-1 combination (Table 3).

**Table 3:** Data structure of analysed Syn-1 combinations with percentage of F_1_ seeds, parental component 1 and parental component 2 to the combinations and tested P-Values from a t-test.

| Synthetics | $F_1$ | P1 | P2 | P-Values Weight | | | P-Values Volume | | |
| --- | --- | --- | --- | --- | --- | --- | --- | --- | --- |
| | [%] | [%] | [%] | $F_1$ - P1 | $F_1$ - P2 | P1 - P2 | $F_1$ - P1 | $F_1$ - P2 | P1 - P2 |
| NPZ-BP3Y2023-001 | 14.6 | 62.5 | 22.9 | 0.7247 | 0.0095 | 0.0006 | 0.8825 | 0.0006 | <0.0001 |
| NPZ-BP3Y2023-006 | 12.5 | 40.6 | 46.9 | 0.6035 | 0.8474 | 0.5839 | 0.0735 | <0.0001 | 0.3866 |
| NPZ-BP3Y2023-007 | 26.9 | 47.3 | 25.8 | 0.6452 | 0.3769 | 0.5794 | 0.0758 | 0.4246 | 0.0027 |
| NPZ-BP3Y2023-020 | 13.5 | 55.2 | 31.3 | 0.1095 | 0.9019 | 0.0246 | 0.0194 | 0.7476 | 0.0001 |
| NPZ-BP3Y2023-021 | 15.8 | 52.6 | 31.6 | 0.0516 | 0.2202 | 0.3456 | 0.1430 | 0.7810 | 0.0998 |
| NPZ-BP3Y2023-023 | 21.9 | 35.4 | 42.7 | 0.0851 | 0.9191 | 0.0345 | 0.0595 | 0.6014 | 0.0718 |
| NPZ-BP3Y2023-036 | 24.5 | 45.7 | 29.8 | 0.4735 | 0.6174 | 0.1390 | 0.7408 | 0.4120 | 0.5230 |
| NPZ-BP3Y2023-047 | 14.7 | 44.2 | 41.1 | 0.0424 | 0.0076 | <.0001 | 0.0274 | <.0001 | 0.0007 |
| NPZ-BP3Y2023-053 | 13.5 | 27.1 | 59.4 | 0.5512 | 0.5883 | 0.8903 | 0.9539 | 0.2587 | 0.0759 |
| NPZ-BP3Y2023-054 | 14.6 | 49.0 | 36.5 | 0.1179 | 0.1019 | 0.8637 | 0.9539 | 0.3093 | 0.1358 |
| NPZ-BP3Y2023-056 | 14.9 | 33.0 | 52.1 | 0.1034 | 0.0128 | <.0001 | 0.0002 | 0.0001 | <.0001 |
| NPZ-BP3Y2023-060 | 24.2 | 23.2 | 52.6 | 0.8321 | 0.3917 | 0.4570 | 0.5252 | 0.1688 | 0.4118 |
| NPZ-BP3Y2023-065 | 14.9 | 39.4 | 45.7 | 0.7637 | 0.3019 | 0.0431 | 0.2402 | 0.9777 | 0.0780 |
| NPZ-BP3Y2023-090 | 19.5 | 24.7 | 55.8 | 0.2128 | 0.3286 | 0.0031 | 0.3020 | 0.1559 | 0.0013 |
| NPZ-BP3Y2023-093 | 21.5 | 47.3 | 31.2 | 0.0430 | 0.9274 | 0.0149 | 0.1072 | 0.9569 | 0.0495 |
| NPZ-BP3Y2023-107 | 18.9 | 37.9 | 43.2 | 0.0570 | 0.6978 | 0.0018 | 0.0659 | 0.4810 | 0.0006 |
| NPZ-BP3Y2023-113 | 21.5 | 66.7 | 11.8 | 0.7392 | <.0001 | <.0001 | 0.8674 | <.0001 | <.0001 |
| NPZ-BP3Y2023-116 | 14.8 | 45.7 | 39.5 | 0.1794 | 0.2671 | 0.0002 | 0.3408 | 0.0903 | 0.0001 |

In some Syn-1 combinations we observed differences in the seed weight and volume from the parental components (Table 3). Only cultivar NPZ-BP3Y2023-056 showed significant differences in seed volume for every comparison between F_1_ seeds and parental components and between the parental components. Otherwise, significant differences were generally observed between parental components but not between of F_1_ seeds and their respective parents.

#### 3.2.2 Differentiation of parental seeds based on spectral patterns

The accuracies of the differentiation of parental seeds based on spectral patterns vary for scenario 1, depending on the dataset and algorithm. The best performing algorithm is SVM with a mean accuracy of 66.8 %. The best performing dataset is dataset 1 (SWP) including spectral, weight, volume and parental information with a mean accuracy of 69.9 %. The weight and volume data does not influence the result significantly. The best performing combination across algorithms and datasets is RF with dataset 1, with a mean accuracy of 90.5 % ranging from 85.4 % up to 95.1 % (Figure 2).

**Figure 2:**
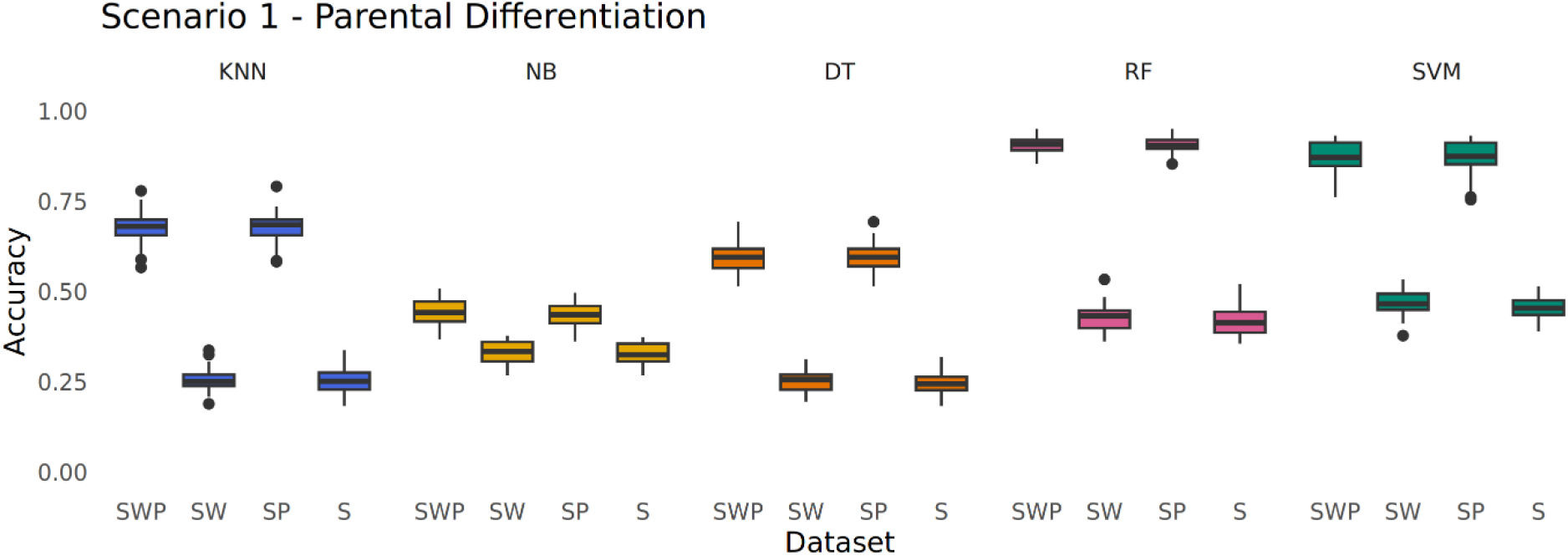
Parental Differentiation of Scenario 1. Comparison of prediction accuracy of differentiation of parental seeds of Syn-1combinations across the different datasets including spectral, weight, volume and parental information (SWP), spectral, weight and volume (SW), spectral and parental information (SP) and only the spectra (S) and different algorithms KNN (blue), NB (yellow), DT (orange), RF (pink) & SVM (green) for scenario 1.

#### 3.2.3 Differentiation of Syn-1 seeds based on spectral patterns

Including all Syn-1 seeds (corresponding to F_1_ seeds and self-pollinated parental components) in the classification tasks reduces the predictability (Figure 3). The F1 score combines precision, reflecting how many of the predicted F_1_ seeds were true F_1_s, and recall, which indicates how many of the actual F_1_ seeds were correctly identified. Overall, the ability to distinguish self-pollinated parental seeds from F_1_ hybrid seeds was poor, as the highest F1 score was only 19.7 %, achieved by the SVM scenario with dataset 1 (SWP). The best mean F1 score across datasets (12.7 %) was obtained via SVM, while and the best dataset was dataset 3 (SP). Inclusion of seed weight and volume did not significantly influence the prediction accuracies (Figure 3a). The most accurate prediction algorithm was SVM with 61.0 %, while the most accurate predictions were achieved in dataset 1 (SWP) with 67.0 %. The best combination is RF with dataset 1 (SWP) with 78.8 % accuracy (Figure 3c). Including the weight and volume in the datasets did not significantly influence the predictions.

**Figure 3:**
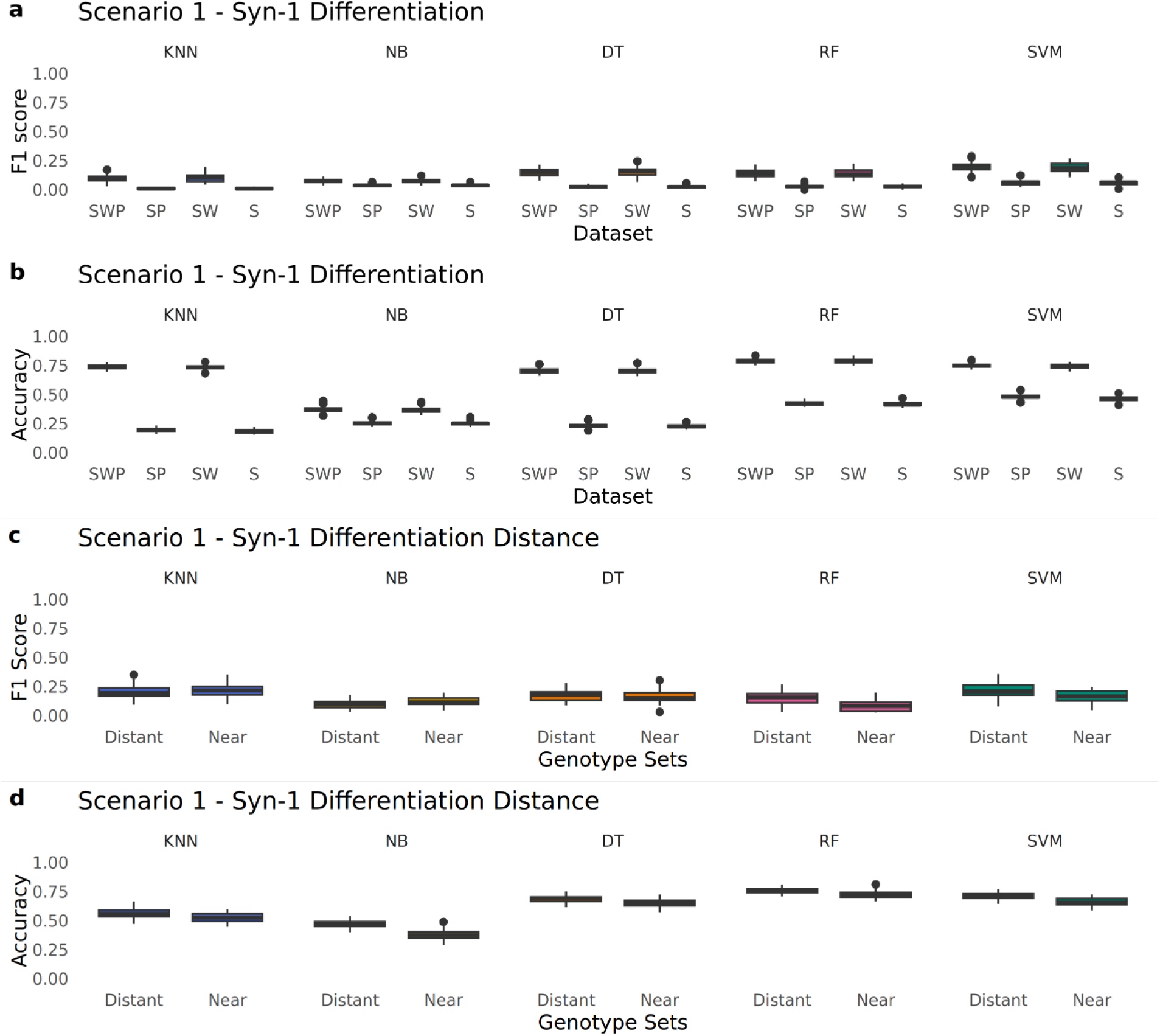
Syn-1 Differentiation of Scenario 1. Comparison of (a) F1 score and (b) prediction accuracy of homozygous/heterozygous components of Syn-1 combinations across the different datasets, including spectral, weight, volume and parental data (SWP, SW, SP &S). (c) F1 score and (d) prediction accuracy for scenario 1 separated into two types of Syn-1 combinations based on PCA in “distant” or “near” Syn-1 combinations. All predications were run utilizing different algorithms KNN (blue), NB (yellow), DT (orange), RF (pink) & SVM (green) for scenario 1.

Separating the Syn-1 combinations into “distant” or “near” groups based on euclidian genetic distance of parental components of Syn-1 combinations, which are already visible in their respective PCAs (Supplementary Figure S1). Superior predictions between “distant” groups for the algorithms DT, RF & SVM for the F1 Score were observed. The best performing algorithm in this respect was KNN with 21.1 %, while the best combination was KNN with “near” genotypes at 21.8 % F1 score (Figure 3b). “Distant” groups of Syn-1 combinations were always predicted better regarding the accuracy, with mean accuracy of 63.8 % in contrast to “near” Syn-1 combinations with 58.7 %. Highest accuracy (75.5 %) was achieved for prediction “distant” Syn-1 combinations with the RF algorithm (Figure 3d).

### 3.3 Stage 2

#### 3.3.1 Data structure

In this stage we analysed the Syn-1 combinations NPZ-BP3Y2023-060 and NPZ-BP3Y2023-007 with a higher number of samples per Syn-1 combination. NPZ-BP3Y2023-060 was represented by a total of 293 seeds, showing an outcrossing rate of 14.72 %, while NPZ-BP3Y2023-007 was represented by 299 seeds and an outcrossing rate of 19.8 %. Differentiation of parental seeds based on spectral patterns Differentiation between the parental components based on the spectral patterns improved in scenario 2 in comparison to scenario 1. The best performing algorithm was SVM with a mean accuracy of 95.6 %. Dataset 4 (S) performs slightly better with 92.7 % accuracy in comparison to dataset 2 (SW) with 92.6 %. The best performing combination across algorithms and datasets was SVM with dataset 2 (SW), with a mean accuracy of 95.6 % ranging from min 88.5 % up to 100.0 % (Figure 4a). Accuracies of differentiation between parental components based on spectral patterns improved in scenario 3 further. The best performing algorithm was RF with a mean accuracy of 98.9 %. Datasets 2 (SW) and 4 (S) performed equally well with 98.6 % accuracy. The best performing combination across algorithms and datasets was RF with datasets 2 (SW) and 4 (S), with a mean accuracy of 98.9 % ranging from min 95.4 % up to 100.0 % (Figure 4b).

**Figure 4:**
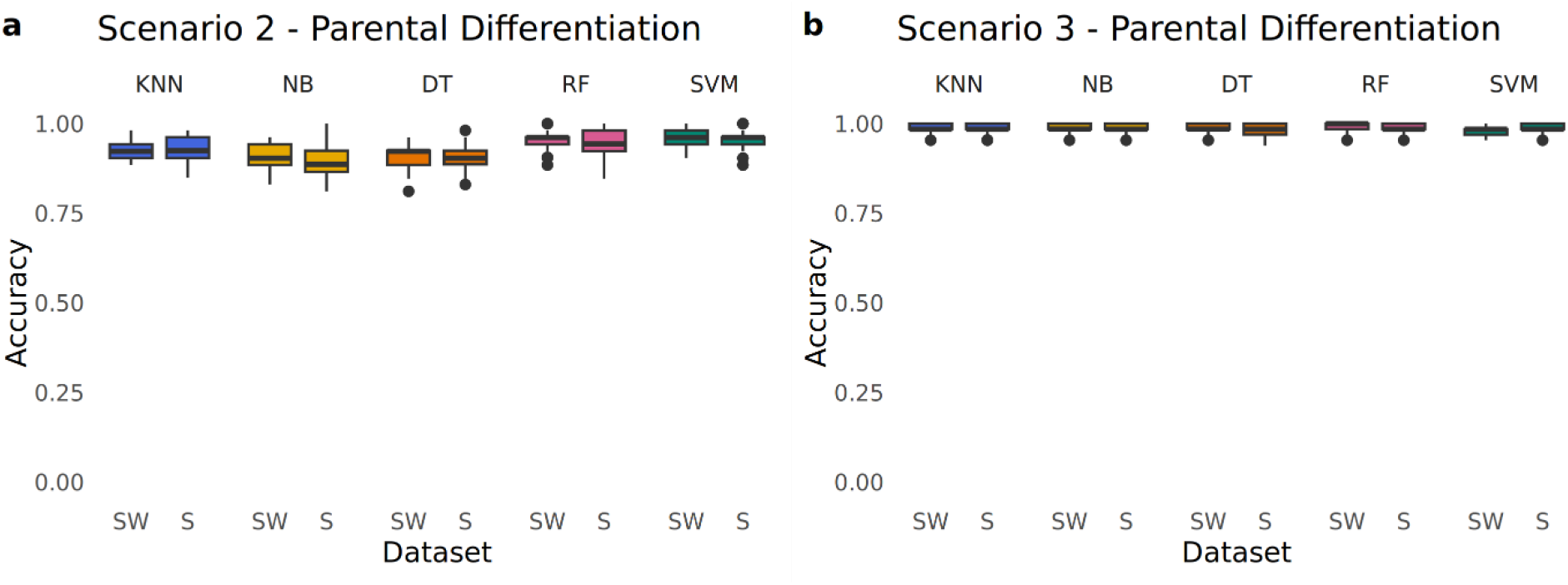
Parental differentiation of Scenario 2 & 3. Comparison of accuracy of differentiation of parental components of Syn-1 combinations using (a) scenario 2 and (b) scenario 3 across the different datasets including spectral, weight and volume data (SW & S) and different algorithms KNN (blue), NB (yellow), DT (orange), RF (pink) & SVM (green).

#### 3.3.2 Differentiation of Syn-1 seeds based on spectral patterns

The accuracy of the differentiation between F_1_ seeds and their parental components based on the spectral patterns was lower than for comparisons between parental components but was improved in scenarios 2 and 3 in comparison to scenario 1. For scenario 2 with Syn-1 combination NPZ-BP32023-060 the F1 score was improved to 38.6 % with RF. The dataset 2 (SW) had the highest mean F1 score with 33.5 %. The best combination was SVM with dataset 2 (SW) and 38.7 % F1 score (Figure 5a). For scenario 3 Syn-1 combination NPZ-BP32023-007 the mean F1 score was 36.8 % with KNN the highest and dataset 2 (SW) and 4 (S) had the highest mean F1 score of 29.2 % while dataset 4 (S) has a higher maximum F1 score of 60.9 %. The best combination was DT with dataset 2 (SW) showing 37.9 % F1 score (Figure 5b). For scenario 2 NPZ-BP32023-060 the accuracy was the highest 61.8 % with RF and dataset 4 (S) with 52.1 %. The best combination with the highest accuracy was RF with dataset 2 (SW) and 4 (S) with 61.9 % (Figure 5c). NPZ-BP32023-007 in the scenario 3 the best algorithm was RF with 60.4 % and the best dataset 2 (SW) with 54.7 %. The best combination between algorithm and dataset was RF and dataset 2 (SW) with 62.8 % (Figure 5d).

**Figure 5:**
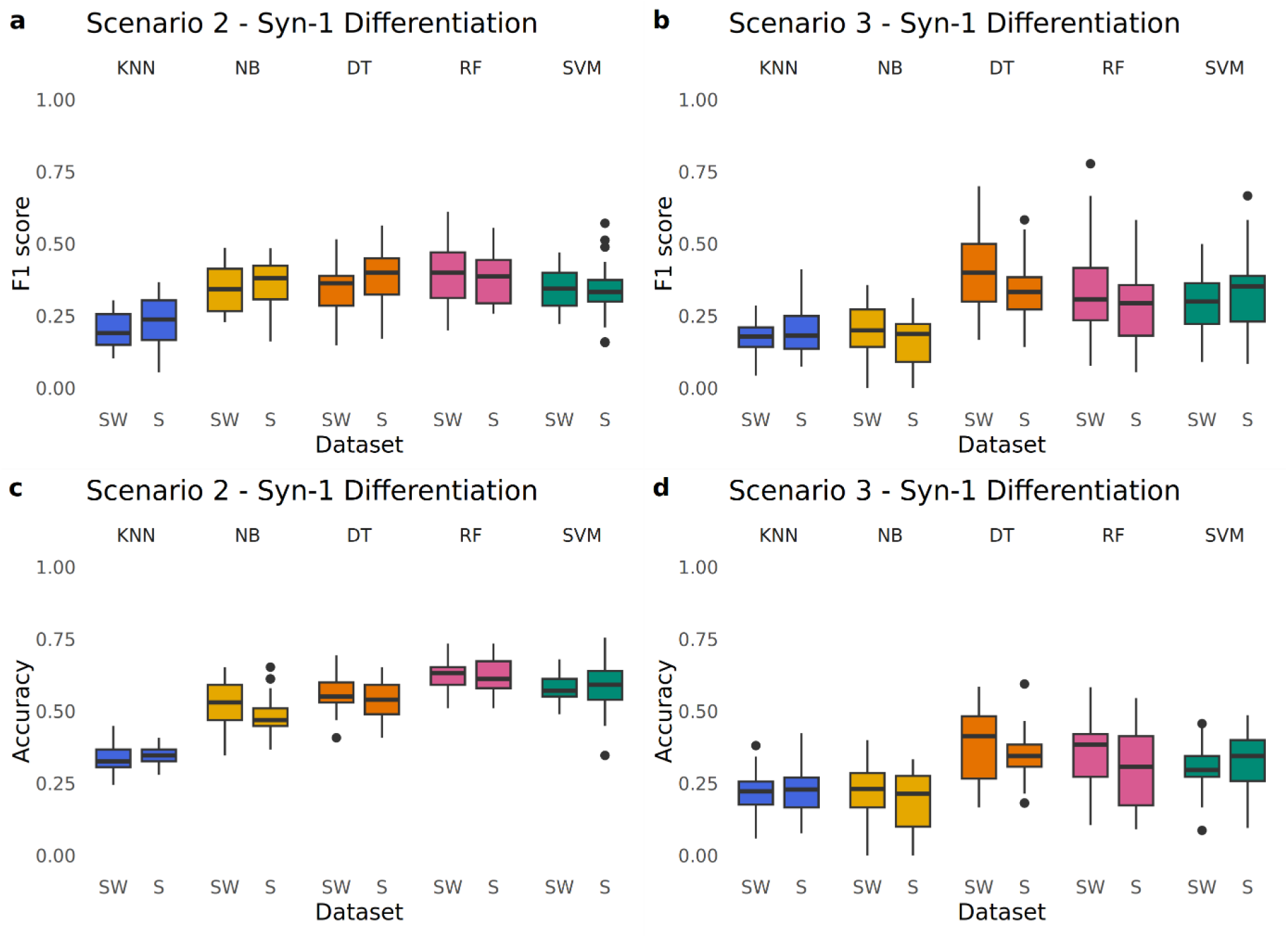
Syn-1 Differentiation of Scenario 2 & 3. Boxplots of prediction of F1 score for scenario 2 (a) and scenario 3 (b) and accuracy for scenario 2 (c) and scenario 3 (d) of differentiation of Syn-1 seeds across the different datasets including spectral, weight and volume (SW & S) and different algorithms KNN (blue), NB (yellow), DT (orange), RF (pink) & SVM (green).

Trying to improve the F1 score the mean spectra was investigated to identify wavelengths showing F_1_ hybrid-specific spectral patterns (Figure 6a & 6b). For scenario 2, algorithm RF performed best in this context with 36.6 % F1 Score and the dataset 2 (SW) with 30.0 %. The best combination for NPZ-BP32023-060 (scenario 2) was RF with dataset 2 (SW) with 36.8 % for the F1 score (Figure 6c). For scenario 3, genotype NPZ-BP32023-007 the best algorithm was SVM with 39.6 % and dataset 2 (SW) with 37.8 %. The best combination of algorithm and dataset was SVM with dataset 2 (SW) with a F1 score of 40.7 % (Figure 6d). The accuracy of scenario 2 is 58.5 % with the algorithm RF, while dataset 2 (SW) showed highest accuracy with SVM (47.5 %). The best combination was DT with dataset 2 (SW) (accuracy 50.4 %). In scenario 3 the highest accuracy between the algorithm was 61.0 % with RF and between datasets, dataset 2 (SW) with a mean accuracy of 59.5 %. The best combination was KNN with dataset 2 (SW) for 61.7 % (Figure 6f).

**Figure 6:**
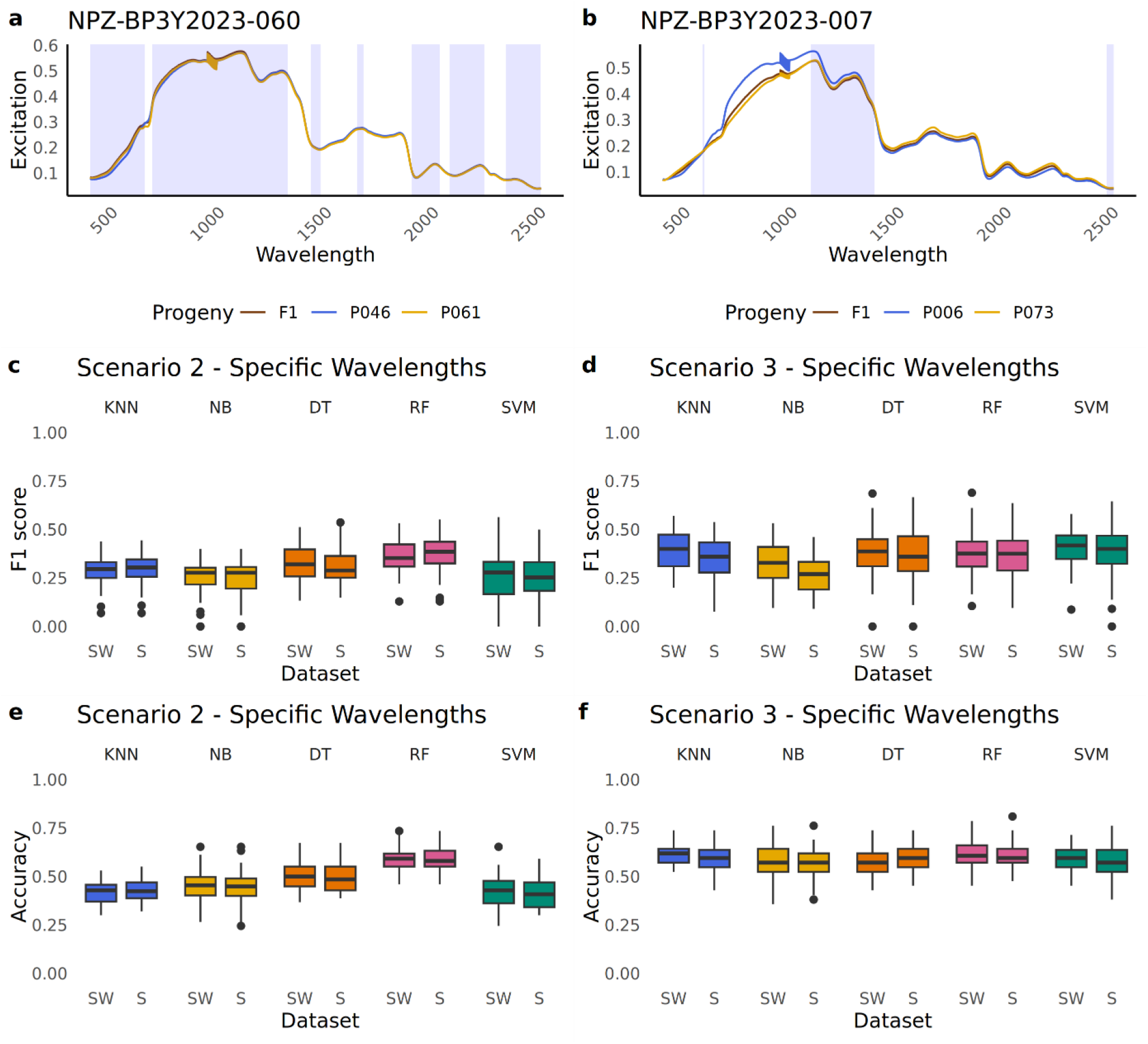
Wavelength specific Syn-1 Differentiation of Scenario 2 & 3. Line graph showing the mean spectra for (a) NPZ-BP3Y2023-060 and (b) NPZ-BP32023-007 generated with the raw spectra, blue background indicating the wavelengths used for the prediction, brown lines for F_1_ seeds and blue and yellow for their parental lines. Boxplots of prediction of F1 score for scenario 2 (c) and scenario 3 (d) and accuracy for scenario 2 (e) and scenario 3 (f) of classification of Syn-1 combinations across the different datasets including spectral, weight and volume (SW & S) and different algorithms KNN (blue), NB (yellow), DT (orange), RF (pink) & SVM (green).

As the parental components of the Syn-1 combination NPZ-BP3Y2023-007 could be separated based on the PCA (Figure 7e). The F_1_ seeds were assigned to one of the parental components, as it is hypothesized that maternal effects lead to this clustering. This can not be validated as we only had a mixture of synthetic combinations and not the exact progeny, which would state the maternal and paternal sides. The parental component P006 of NPZ-BP3Y2023-007 had an approximate outcrossing rate of 13.29 %. The best performing algorithm regarding the F1 score was DT with 62.2 % and dataset 4 (S) with 50.8 %. The best combination was DT with dataset 2 (SW) with 62.2 % for the F1 score (Figure 7a) The accuracy was with DT with 64.6 % the highest and dataset 4 (S) with 53.9 %. The best combination between datasets and algorithm was DT and dataset 4 (S) with the prediction accuracy of 64.8 % (Figure 7c). The parental component P073 had an approximate outcrossing rate of 27.61 %. The F1 score was the highest with the algorithm NB of 60.2 % and the dataset 2 (SW) of 43.4 %. The best combination was NB with dataset 4 (S) with 60.4 % F1 score (Figure 7b). The prediction accuracy was the highest as well for NB with 58.8 % and dataset 2 (SW) with 43.7 %. The best combination with the highest accuracy was NB with dataset 4 (S) 59.0 % (Figure 7d).

**Figure 7:**
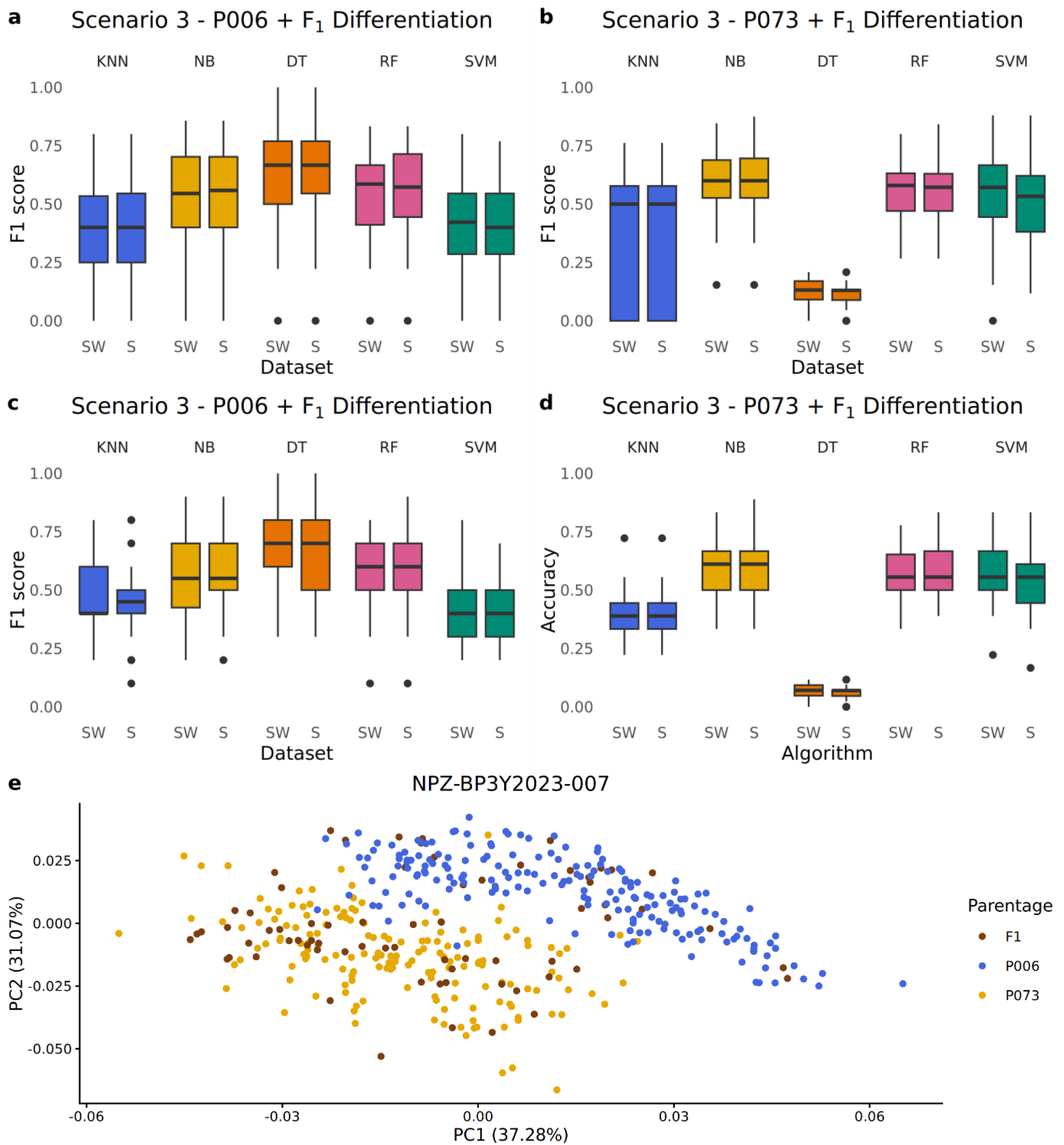
Parental specific Syn-1 differentiation of scenario 3. (a-d) Boxplots describing the ability to differentiate F_1_ seeds from their parents using scenario 3, showing the F1 scores for (a) parental component P006 and (b) parental component P073, as well as the accuracy for (c) parental component P006 and (d) parental component P073 across different datasets including hyperspectral reflectance, weight and volume (SW & S) and different algorithms KNN (blue), NB (yellow), DT (orange), RF (pink) & SVM (green). (e) PCA of hyperspectral reflectance patterns from seeds representing F_1_ seeds (brown points) and their respective parents (blue and yellow points, respectively) from the biparental Syn-1 combination NPZ-BP3Y2023-007.

## 4 Discussion

### 4.1 Identification of parental components of Syn-1 combinations

Numerous studies have addressed the assessment of variety purity using spectral data. For instance, variety identification based on spectral analysis has been extensively documented in crops such as wheat, maize, okra, and loofah seeds (Zhao et al. 2022b, Yang et al. 2015, Nie et al. 2019). These studies often integrate spectral and spatial features, achieving classification accuracies of up to 98 %. In the present study, stage 1 demonstrated that, despite the limited sample size per group, the parental components of faba bean Syn-1 combinations could be classified with an accuracy of 90.5 %. The scenarios developed in stage 2 further enhanced predictive performance, reaching accuracies of up to 98.9 %. Notably, while Syn-1 combinations differences were observed, prediction accuracies remained consistently high.

### 4.2 Are xenia effects in faba bean revealed by spectral patterns?

The xenia effect in faba bean seeds was described by Duc et al. (2001), who observed an influence of the paternal pollen on the volume and number of cotyledon cells for all analysed faba bean genotypes. However, the impact on seed weight was limited to small-seeded paternal components. In our study, no significant differences were detected between the parental components and the F_1_ seeds across the 18 Syn-1 combinations examined, except for Syn-1 combination NPZ-BP3Y2023-056 (Table 3). For this Syn-1 combination, the volume differed significantly not only between the parental components and the F1 seeds but also between the parental components themselves. Heterosis for F1 seed weight, particularly when parents are genetically divergent, has been described by Dieckmann and Link (2010). While such positive effects on seed weight are documented in hybrid seeds, they were not reflected in the predictive performance of our scenarios. Neither seed weight nor volume exhibited a significant influence on our prediction performance. In different species like maize an increased seed weight in hybrid seed is observed as well, and is considered an early form of heterosis (Bulant and Gallais 1998).

A xenia effect on seed colour, as observed in pea with interspecific crosses (Sari et al. 2023), has not yet been reported in faba bean, where seed colour is determined by maternal inheritance (Ricciardi et al. 1985). This is consistent with soybean, in which seed colour has also been described as maternally inherited (Kumar et al. 2025). These observations are supported by the clustering patterns in PCAs, where in parental components formed two distinct clusters and F_1_ seeds fell into one or other of the parental clusters. Under these conditions the expected outcome would be 33 % correct classifications through random assignment in stage 2 of this study for the F1 score. However, our scenario achieved a mean F1 score of 39 %, indicating that a xenia effect may be present but either minimal or not detectable within the visible spectrum. This is further supported by the differentiation of Syn-1 combinations with separated parental components, where random assignment would have an accuracy or F1 score of 50 %, yet our algorithms exceeded this threshold, reaching up to 64 % accuracy. To improve predictive performance, a potential next step could involve the development of a hybrid convolutional neural network designed to capture both the spectral reflectance and the structural or textural features of seeds. Zhao et al. (2022b) successfully applied this approach to wheat variety identification, demonstrating that a model combining spectral and spatial features outperformed those based on either feature set alone.

### 4.3 Faba bean breeding applications

Faba bean are bred as synthetic cultivars, with a mean outcrossing rate of approximately 50 % and a range from 7 % to 82 % (Link 1990, Link et al. 1994a, Link et al. 1994b). An ability to distinguish F_1_ seeds from their homozygous parents could potentially be used to enrich F_1_ hybrid seeds in synthetic cultivars, thus potentially improving heterosis and seed yield (Link et al. 1994b). Moreover, the technique might also be useful as a selection method to identify parental components that show elevated outcrossing rates.

Considering the outcrossing rates of the Syn-1 combinations used in these studies, which ranged from 13.5 % to 26.9 %, an F1 score of 39 %, like in scenario 2, would correspond to a hybrid proportion of 39 % in the sorted cultivar. Despite the relatively modest prediction performance, this would still be expected to result in a yield increase, as previously described. Furthermore, such predictions or sorting could potentially reduce the number of propagation generations required for synthetic cultivars. Typically, Syn-2 or Syn-3 generations are sold to farmers (Stelling et al. 1994). If the desired yield level could be achieved as early as the Syn-1 generation through seed sorting, this could save up to one generation of costly seed propagation.

Overall, while this method has not yet led to the implementation of a 100% pure hybrid cultivar, it could pave the way for hybrid-enriched synthetic cultivars and demonstrate the potential of using hyperspectral images for seed sorting.

## 5 Conclusion

In conclusion, the differentiation of parental components in Syn-1 combinations from biparental synthetic combinations via spectral patterns generally performs well. While the accuracy of the identification of F_1_ seeds is comparatively lower, it is nevertheless sufficient for potential enrichment of Syn-1 seedlots with F_1_ hybrid seeds. This sorting approach could result in an increased proportion of F_1_ hybrid seeds within synthetic cultivars, potentially boosting yield due to heterosis. Better performance of Syn-1 cultivars compared to Syn-2 or Syn-3 would reduce the number of propagation cycles required, saving time and costs in breeding programmes for synthetic cultivars. This study demonstrates the potential of seed hyperspectral reflectance data and machine learning to improve in faba bean breeding. Implementation in conjunction with automated seed sorting could further streamline breeding processes and contribute to possible time, cost and yield advantages in development of synthetic cultivars.

## 6 Acknowledgement

The authors gratefully acknowledge Sven Ernst Weber and Lennart Roscher-Ehrig, Department of Plant Breeding Justus Liebig University Giessen, for providing valuable feedback and supervision.

## 7 Author Contribution Statement

**Rica-Hanna Schlichtermann:** Conceptualization, Methodology, Formal analysis, Investigation, Writing – Original Draft. **Sebastian Warnemünde:** Methodology, Formal analysis, Investigation. **Hanna Tietgen:** Resources. **Gregor Welna:** Resources. **Andreas Stahl:** Supervision. **Benjamin Wittkop:** Conceptualization, Supervision. **Rod J. Snowdon:** Conceptualization, Supervision, Funding acquisition. All authors revised the manuscript and approved the final manuscript.

## 8 Data availability

The dataset generated during the current study is available under reasonable request.

## 11 Declaration of generative AI and AI-assisted technologies in the writing process

During the preparation of this work the lead author used Le Chat from Mistral AI (Knowledge cutoff: November 2024) in order to improve language and readability. After using this tool, the authors thoroughly reviewed and edited the content as needed. The authors take full responsibility for the content of the published article.

## 13 Supplementary Materials

**Figure S1:**
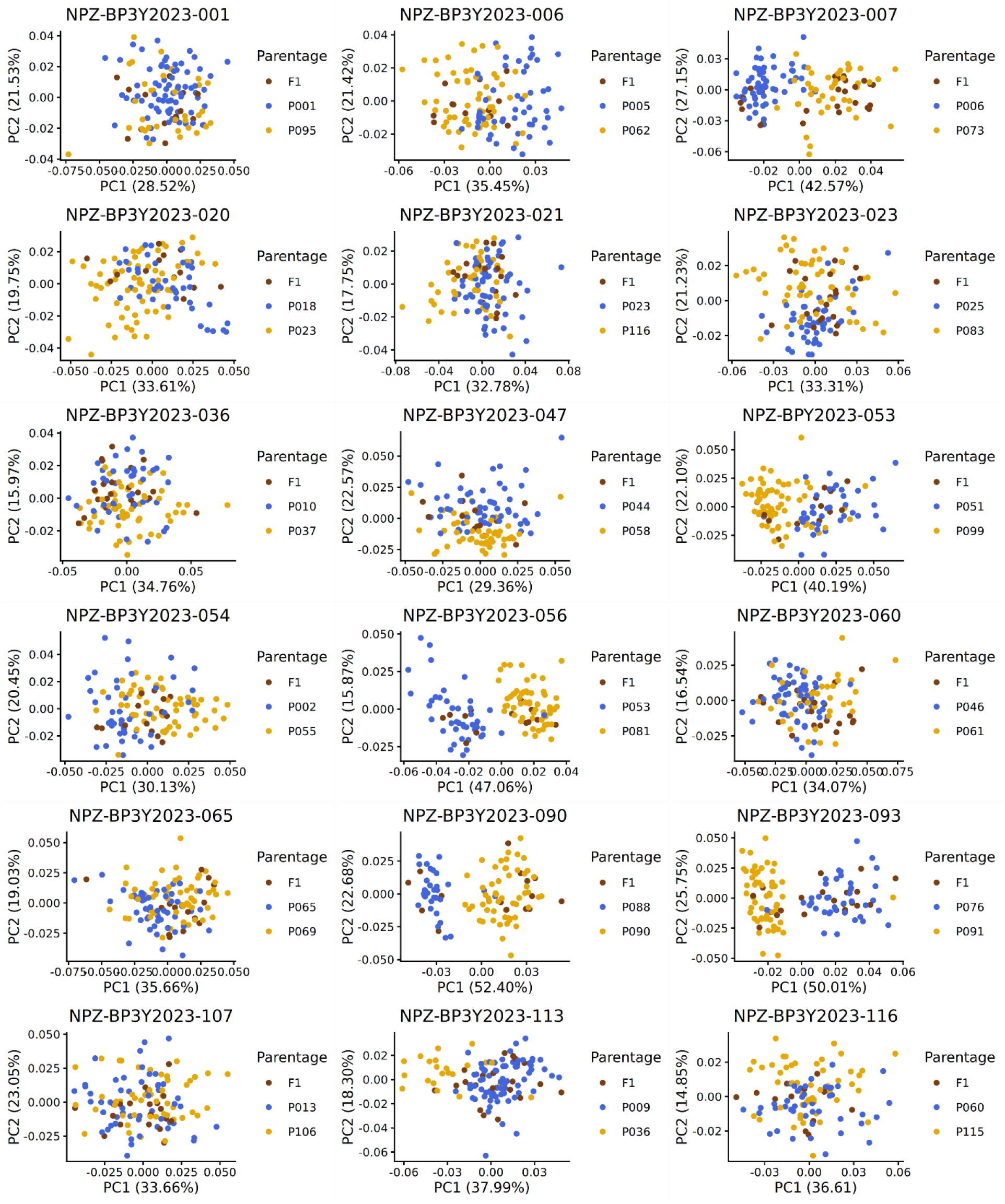
Principal component analysis of hyperspectral reflectance patterns of Syn-1 combinations. PCA of hyperspectral reflectance patterns from seeds representing F_1_ hybrids (brown points) and their respective parents (blue and yellow points, respectively) from 18 biparental Syn-1 synthetic faba bean combinations.

**Figure S2:**
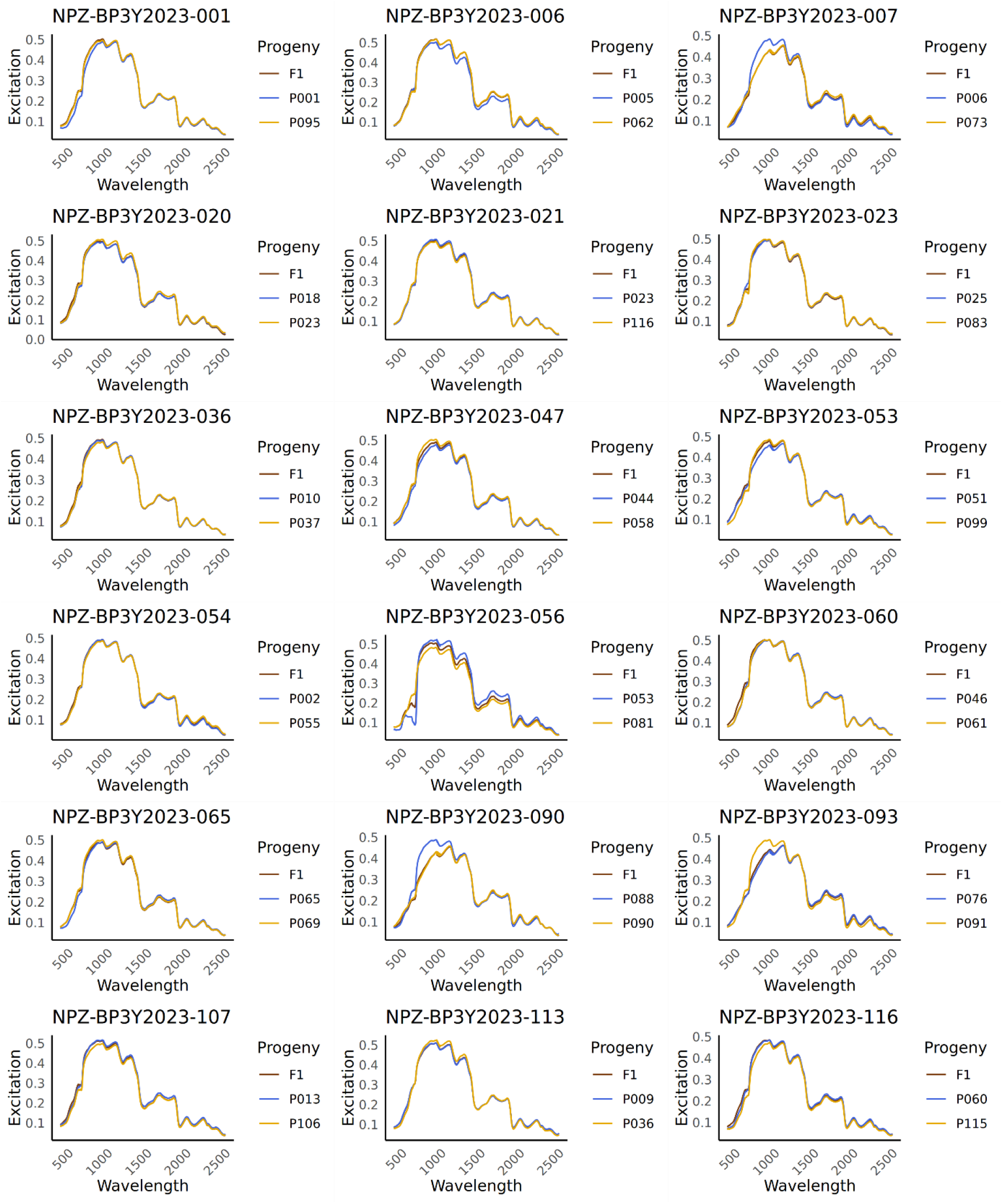
Mean hyperspectral reflectance patterns of Syn-1 combinations. Mean hyperspectral reflectance patterns from seeds representing F_1_ hybrids (brown lines) and their respective parents (blue and yellow lines, respectively) from 18 biparental Syn-1 synthetic faba bean combinations based on the raw spectra.

